# Dioxin Disrupts Thyroid Hormone and Glucocorticoid Induction of *klf9*, a Master Regulator of Frog Metamorphosis

**DOI:** 10.1101/2021.12.01.470809

**Authors:** David T. Han, Weichen Zhao, Wade H. Powell

**Affiliations:** Biology Department Kenyon College Gambier, OH 43022 USA

**Keywords:** Endocrine disruption, aryl hydrocarbon receptor, metamorphosis, dioxin, thyroid, glucocorticoid

## Abstract

Frog metamorphosis, the development of an air-breathing froglet from an aquatic tadpole, is controlled by thyroid hormone (TH) and glucocorticoids (GC). Metamorphosis is susceptible to disruption by 2,3,7,8-tetrachlorodibenzo-*p*-dioxin (TCDD), an aryl hydrocarbon receptor (AHR) agonist. Krüppel-Like Factor 9 (*klf9*), an immediate early gene in the endocrine-controlled cascade of expression changes governing metamorphosis, can be synergistically induced by both hormones. This process is mediated by an upstream enhancer cluster, the *klf9* synergy module (KSM). *klf9* is also an AHR target. We measured *klf9* mRNA following exposures to triiodothyronine (T3), corticosterone (CORT), and TCDD in the *Xenopus laevis* cell line XLK-WG. *klf9* was induced 6-fold by 50 nM T3, 4-fold by 100 nM CORT, and 3-fold by 175 nM TCDD. Co-treatments of CORT and TCDD or T3 and TCDD induced *klf9* 7- and 11-fold, respectively, while treatment with all 3 agents induced a 15-fold increase. Transactivation assays examined enhancers from the *Xenopus tropicalis klf9* upstream region. KSM-containing segments mediated a strong T3 response and a larger T3/CORT response, while induction by TCDD was mediated by a region ∼1 kb farther upstream containing 5 AHR response elements (AHREs). This region also supported a CORT response in the absence of readily-identifiable glucocorticoid responsive elements, suggesting mediation by protein-protein interactions. A functional AHRE cluster is positionally conserved in the human genome, and *klf9* was induced by TCDD and TH in HepG2 cells. These results indicate that AHR binding to upstream AHREs represents an early key event in TCDD’s disruption of endocrine-regulated *klf9* expression and metamorphosis.

## Introduction

Frog metamorphosis, the developmental transition from an aquatic tadpole to a terrestrial froglet, involves substantial, widespread tissue remodeling, including tail resorption, limb growth, intestinal shortening, alterations to the central nervous system, and conversion of the respiratory system to enable air breathing (reviewed throughout Shi (2000)). Metamorphosis starts and proceeds under endocrine control, mediated primarily by thyroid hormones (TH) with substantial modulation by glucocorticoids (GC; Sachs and Buchholz, 2019). The tractable physical, chemical, and genetic manipulation of *Xenopus* embryos, larvae, and tadpoles (*X. laevis* and *X. tropicalis*) has enabled their emergence as a model system for examining the basic mechanisms of endocrine control of development and its disruption by xenobiotics to yield adverse outcomes, particularly the TH- and GC-controlled events supporting metamorphosis (*e.g.*, US EPA, 2011; Helbing *et al*., 2007a; Helbing *et al*., 2007b; Mengeling *et al*., 2017; Miyata and Ose, 2012; OECD, 2009; Thambirajah *et al*., 2019).

Dioxin-like chemicals (DLCs), including chlorinated dioxins and furans and planar polychlorinated biphenyls, are environmental contaminants that can disrupt the endocrine control of metamorphosis in *Xenopus* (Taft *et al*., 2018). 2,3,7,8 tetrachlorodibenzo-*p*-dioxin (TCDD) exposure speeds resorption of cultured tail explants while slowing tadpole limb growth. These morphological effects are likely related to the altered expression of several genes that drive the earliest molecular events in metamorphosis, including *klf9*, which is also a target of both TH and GC (Taft, et al., 2018). The effects of TCDD and other DLCs are mediated by the aryl hydrocarbon receptor (AHR), a ligand-activated transcription factor. DLC agonists bind the cytoplasmic receptor, triggering its translocation to the nucleus, where it forms a heterodimer with the ARNT protein and binds cognate enhancer elements (AHREs, also known as XREs or DREs), affecting the transcription of target genes (reviewed in Gasiewicz and Henry, 2012).

TH and GC also exert their effects through specific binding to their respective nuclear receptors. In the context of tadpole metamorphosis, each hormone directly or indirectly affects hundreds of tadpole genes individually or in concert (Kulkarni and Buchholz, 2012). The most potent form of TH, T3 (triiodothyronine; converted by deiodinases from thyroxine (T4) at target tissues (Bianco *et al*., 2019)), initially binds the constitutively expressed thyroid receptor alpha (TRα) and subsequently the T3-induced TRβ. Each TR can form a functional dimeric transcription factor in concert with the retinoid X receptor (RXR), binding any of several sequences that define a thyroid- responsive element (TRE). The TRα:RXR heterodimer binds constitutively to DNA, acting as a transcriptional repressor in the unbound state and an activator following T3 binding. These “dual functions” serve to regulate the timing and inducibility of metamorphic events, inhibiting them prior to TH production and stimulating them thereafter as a result of receptor binding (reviewed in Buchholz *et al*., 2006; Buchholz and Shi, 2018). Glucocorticoids, widely associated with the stress response, play important roles in vertebrate development, often speeding processes that define transitions between life stages (Wada, 2008). These adrenal corticosteroids, which include corticosterone and cortisol, act by binding the glucocorticoid receptor (GR).

Ligand binding triggers GR translocation from the cytoplasm to the nucleus, where it interacts directly with DNA (classically as a homodimer) at a glucocorticoid response element (GRE) or through interactions mediated by additional transcription factors or cofactors (reviewed in Scheschowitsch *et al*., 2017).

The *klf9* gene, encoding the basic leucine zipper protein krüppel-like factor 9 (a.k.a. “BTEB”), is among the earliest targets for transcriptional induction by TH and GC in the cascade of molecular events that results in metamorphosis. Klf9 protein in turn acts as a key regulator of subsequent transcriptional changes that drive this process (Furlow and Kanamori, 2002). Recent studies have highlighted the synergistic nature of *klf9* induction by TH and GC, identifying a ∼180-bp cluster of enhancers known as the “*klf9* synergy module” (KSM) that regulates *klf9* induction. The KSM, well conserved among tetrapods, contains functional versions of both GRE and TRE motifs (Bagamasbad *et al*., 2015).

Our own previous studies in multiple *X. laevis* cell lines and whole tadpoles demonstrate that *klf9* mRNA expression is induced directly by TCDD in AHR-dependent fashion, while co-exposure to TCDD and T3 can boost *klf9* expression in apparently synergistic fashion (Taft, et al., 2018). This phenomenon likely contributes substantially to the morphological perturbations TCDD elicits in developing tadpole tissues. In the present study, we probed the mechanism by which TCDD disrupts the endocrine control of *klf9* expression, seeking to identify enhancers that mediate the role of AHR and their relationship with the KSM in the altered transcriptional response to both T3 and corticosterone. We report the identification of a functional cluster of AHREs in both the frog and human genomes as well as a region upstream from the KSM that supports GC- induced *klf9* transcription.

## Materials and Methods

### Chemicals

3,3’,5-Triiodo-L-thyronine sodium salt (T_3_), corticosterone (CORT), and 2,3,7,8- Tetrachlorodibenzo-*p*-dioxin (TCDD) were purchased from Sigma-Aldrich (St. Louis, MO). Solutions of T_3_, CORT, and TCDD were prepared in dimethyl sulfoxide (DMSO) obtained from Sigma Aldrich. MS-222 was obtained from Western Chemical Company (Ferndale, WA).

### Cell Culture

#### Frog cells

The *Xenopus laevis* adult renal epithelial cell line, XLK-WG (ATCC; Manassas, VA), is a longstanding and well-characterized frog cell line for studying cellular responsiveness to AHR agonists (Laub *et al*., 2010; Freeburg *et al*., 2016; Iwamoto *et al*., 2012; Taft, et al., 2018). XLK-WG cells were cultured as directed by the supplier at 29°C/5% CO_2_ in RPMI-1640 medium supplemented with 20% fetal bovine serum (FBS; ThermoFisher) and passaged at ∼70% confluence using a 0.25% trypsin EDTA solution in 1X PBS. Media were used and stored with care to minimize formation of 6-formylindolo[3,2*b*]carbazole (FICZ) and related AHR agonists resulting from light exposure (Oberg *et al*., 2005).

#### Human cells

The HepG2 cell line (ATCC) was used to interrogate *klf9* induction by TCDD and T3 cotreatment. Cells were maintained as directed by the supplier in ATCC-formulated Eagle’s Minimum Essential Medium (EMEM) supplemented with 10% FBS. *Exposures and RNA purification.* For experiments measuring *klf9* mRNA in *X. laevis*, XLK-WG cells were cultured to 70% confluence in T-25 flasks. To facilitate direct comparison, cells were treated as described in previously published studies with 100 nM CORT, 50 nM T_3_ (high hormone concentrations to maximize response; Bonett *et al*., 2010), 175 nM TCDD (Taft, et al., 2018), or combinations thereof for 24 hours. This high concentration of TCDD represents the lower-bound EC50 for cyp1a6 induction in this cell line (Laub, et al., 2010), reflecting the relatively low affinity of frog AHRs for TCDD (Lavine et al. 2005). All exposure groups, including controls, held the concentration of DMSO vehicle at 0.25% in RPMI-1640 supplemented with 10% charcoal-stripped FBS. Similarly, concentrations of exposure agents were chosen for HepG2 cells to facilitate direct comparisons with previously published work, using high concentrations to maximize the response. Cells were exposed as described previously to TCDD (10 nM; Dere *et al*., 2011; Jennen *et al*., 2011), T3 (100 nM; Cvoro *et al*., 2015) and/or DMSO vehicle (0.25%) for 24 hours.

### In vivo experiments

*Animals*. *Xenopus laevis* tadpoles (NF 52-54; Nieuwkoop and Faber, 1994) were obtained from Nasco (Fort Atkinson, WI). Protocols for tadpole experiments were approved by the Kenyon College Institutional Animal Care and Use Committee. *Exposures*. During exposures, tadpoles were kept at a density of 3 individuals per 250 ml FETAX solution, a widely used medium containing dilute ions that is optimized for toxicological studies involving *Xenopus* embryos and tadpoles (ASTM, 2012; Dawson and Bantle, 1987; Taft, et al., 2018), under a 12L:12D photoperiod at 23°C. To facilitate direct comparison, tadpoles were treated as described for previously published studies with 100 nM CORT, 50 nM T_3_ (high hormone concentrations to maximize response; Bonett *et al*., 2010), 50 nM TCDD (Taft, et al., 2018), or combinations thereof for 24 hours. All exposures, including controls, held the concentration of DMSO vehicle at 0.25%. At the end of the exposure period, tadpoles were sacrificed by rapid anesthetic overdose in bicarbonate-buffered MS-222 (0.2%) and immediately flash-frozen in liquid nitrogen.

### *klf9* Enhancer Analysis

#### Reporter plasmids

To determine the positions of response element sequences relative to *klf9* in *X. tropicalis* and human genomes, we obtained their respective upstream sequences from the Xenbase genome browser (*X. tropicalis* v9.1; Karimi *et al*., 2018) or the UCSC human genome browser (GRCh38/hg38; Kent *et al*., 2002). Conserved aryl hydrocarbon response elements, glucocorticoid response elements, and thyroid hormone response elements within 10 kb of the annotated transcription start site were identified by transcription factor subsequence search in MacVector 18.1, which includes the tfsites collection (http://www.ifti.org) as well as additional curated sequences. This effort was augmented by manual searches.

Dr. Pia Bagamasbad and Dr. Robert Denver (University of Michigan) generously provided plasmid constructs spanning 1kb intervals up to position -7000 of *X. tropicali*s *klf9* in the pGL4.23 firefly luciferase reporter vector. We designed additional constructs containing human or *X. tropicalis* sequences (indicated in the figures) for synthesis by Epoch Life Sciences (Sugar Land, TX). Reporter plasmids were transformed into chemically competent JM109 *E. coli* (Promega, Madison, WI) and isolated in large quantities using a ZymoPURE-EndoZero Maxiprep kit (Zymo Research, Irvine, CA) or a HiSpeed Plasmid Maxi Kit (Qiagen). To examine the functionality of select putative AHR or GR binding sites, elements were mutagenized to abrogate receptor binding in the pGL4.23 construct containing elements from -5.7 to -7.7 kb. Mutant constructs containing point mutations of 5 putative AHREs (singly or collectively; 5’CACGC3’ > 5’CATGC3’) (Karchner *et al*., 1999; Powell *et al*., 1999) and 1 GRE (5’AGAACAGT3’ > 5’TAGCATCT3’) were generated by site-directed mutagenesis (Epoch Life Sciences). *Transactivation Assays*. To characterize putative response elements of the *X. tropicalis klf9* upstream region, we conducted a series of transactivation assays. We seeded 48- well plates with XLK-WG cells at a density of 10,000 or 15,000 cells per well prior to overnight incubation. Each well was then transfected with 188-220 ng of a pGL4.23 firefly luciferase reporter vector containing a segment of the *X. tropicalis klf9* upstream region. Cells were also co-transfected with 15 ng pTK-*Renilla* (Promega) plasmids using Lipofectamine 3000 reagent (Thermo Fisher Scientific). Cells were incubated for 24 to 48 hours in the transfection solution. Following transfection, populations of plated cells were dosed with DMSO vehicle (0.25%), CORT, T_3_, and/or TCDD for 24 hours as indicated in the figure legends. Reporter gene expression was assessed in 20 µl of cell lysates using the Dual Luciferase Assay and a GloMax luminometer (Promega). To control for transfection efficiency, the degree of transactivation was expressed as relative luciferase units (RLU), the ratio of reporter-driven firefly luciferase activity to constitutive *Renilla* luciferase activity (driven by pTK-*Renilla*). The mean RLU value for each treatment group was calculated for each plasmid relative to the corresponding vehicle control, resulting in the plotted fold change values. At least three biological replicates were conducted for each experiment, each with three repeated measures.

Assays in HepG2 cells were performed similarly to examine *KLF9* enhancers in the human genome.

Data were plotted and analyzed statistically using Prism 8.4.3 (GraphPad). Dual luciferase assay data were log-transformed prior to statistical analysis, facilitating direct comparison with a previously-published workflow for *klf9* enhancer analysis (Bagamasbad, et al., 2015). We followed a similar protocol for our HepG2 cell line transactivation assays, modifying only the exposure concentrations and media as described above.

### mRNA expression analysis

#### RNA purification from cultured cells

Total RNA was purified using QIAshredder and RNeasy kits (Qiagen), using on-column DNase treatment as directed to remove residual genomic DNA.

#### RNA purification from tadpoles

Whole X. *laevis* tadpoles (NF 52-54) were homogenized in RNA STAT-60 (Tel-Test, Inc., Friendswood, TX) and total RNA purified using a Direct-zol RNA miniprep kit (Zymo Research). We used an RNA Clean and Concentrator Kit (Zymo Research) with RNase-Free DNase (QIAGEN, Germantown, MD) and/or DNAfree TURBO (ThermoFisher) to remove residual genomic DNA. *Quantitative RT-PCR*. mRNA measurements were performed essentially as described previously (Freeburg, et al., 2016; Laub, et al., 2010; Taft, et al., 2018; Zimmermann *et al*., 2008). For both *X. laevis* and human RNA, cDNA synthesis reactions were performed using the TaqMan Reverse Transcription Reagent kit (Thermo Fisher Scientific) with random hexamer primers. Reaction conditions were as prescribed by the manufacturer: 25°C/10 m, 37°C/30 m, 95°C/10 m. Quantitative real-time PCR was performed using PowerupSYBR Green 2X PCR Master Mix (Thermo Fisher Scientific) and oligonucleotide primers (Eurofins Genomics, Louisville, KY; Tables 1 and 2) on a 7500 Real-Time PCR system (Thermo Fisher Scientific). For *X. laevis* tadpole and XLK- WG samples, induction of the target gene *klf9* (*klf9.L*) was quantified relative to expression of the *actb* endogenous control (Freeburg, et al., 2016; Laub, et al., 2010; Taft, et al., 2018; Zimmermann, et al., 2008). For human cell line experiments, we measured the relative amplification of *KLF9* and *CYP1A1* using the widely employed endogenous control *GAPDH*. Cycling parameters: 95 °C/10ʹʹ; [95 °C/15ʹʹ, 60 °C/60ʹʹ] x 40 cycles.) To verify the amplification of a single target, melting point analysis was performed after each reaction. Target gene induction was quantified by the ΔΔCt method using SDS 1.4 software (Thermo Fisher Scientific). Data were plotted and statistical analyses were conducted using Prism 8.4.3 (GraphPad).

**Table 1:**
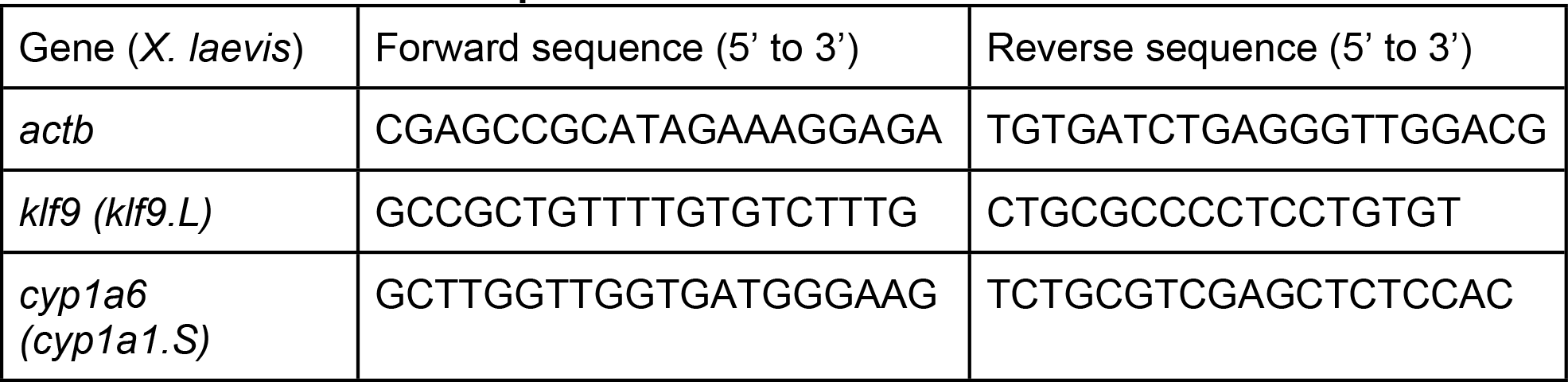
RT-PCR Primer sequences used with XLK-WG cells.

**Table 2.**
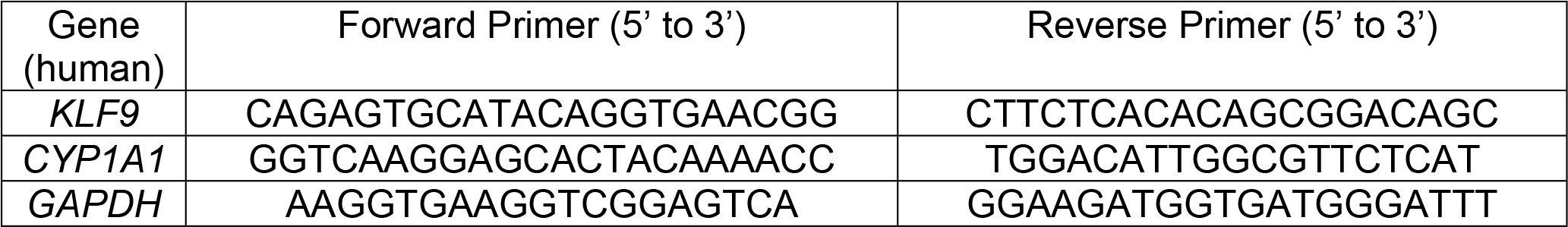
RT-PCR Primer sequences used with HepG2 cells.

## Results

### *klf9* is induced by thyroid hormone, corticosterone, and TCDD

Previously published work characterized the combined effects of T3 and CORT (Bagamasbad, et al., 2015) or T3 and TCDD (Taft, et al., 2018) on *klf9* induction in *Xenopus*. This study extends previous studies by exploring the mechanisms underlying the intersecting functions of both hormones and the AHR pathway, which have not been simultaneously examined. To probe the interactions of T3, CORT and TCDD in altering *klf9* expression in the frog model, we used the XLK-WG cell line, the most extensively-characterized model of frog AHR signaling (Laub, et al., 2010; Freeburg, et al., 2016; Iwamoto, et al., 2012; Taft, et al., 2018), derived from *X. laevis* renal epithelium.

Relative to vehicle control, XLK-WG *klf9* mRNA expression was elevated 6-fold by 50 nM T3, 4-fold by 100 nM CORT, and 3-fold by 175 nM TCDD following 24-hour exposures (Fig. 1). Combined exposures drove further increases in *klf9* induction. Co-treatment with T3 and TCDD induced *klf9* mRNA 11-fold, while CORT and TCDD together yielded a 7-fold relative increase. Simultaneous exposure to T3, CORT, and TCDD increased expression 16-fold (Fig. 1). A similar induction pattern was observed for *klf9* mRNA in whole tadpoles (NF 52-54) exposed for 24 hours to 50 nM T3, 100 nM CORT, and 50 nM TCDD, individually or in combination. Mean *klf9* mRNA displayed apparent increases of 2- to 5-fold to single agents, and exposure to pairwise combinations (CORT + TCDD or CORT + T3) boosted expression further. Coexposure to all three compounds did not elicit an increase greater than either pairwise combination (Fig. 2). Nonetheless, each of the three chemicals was in some fashion associated with elevated *klf9* mRNA *in vivo*.

**Figure 1.**
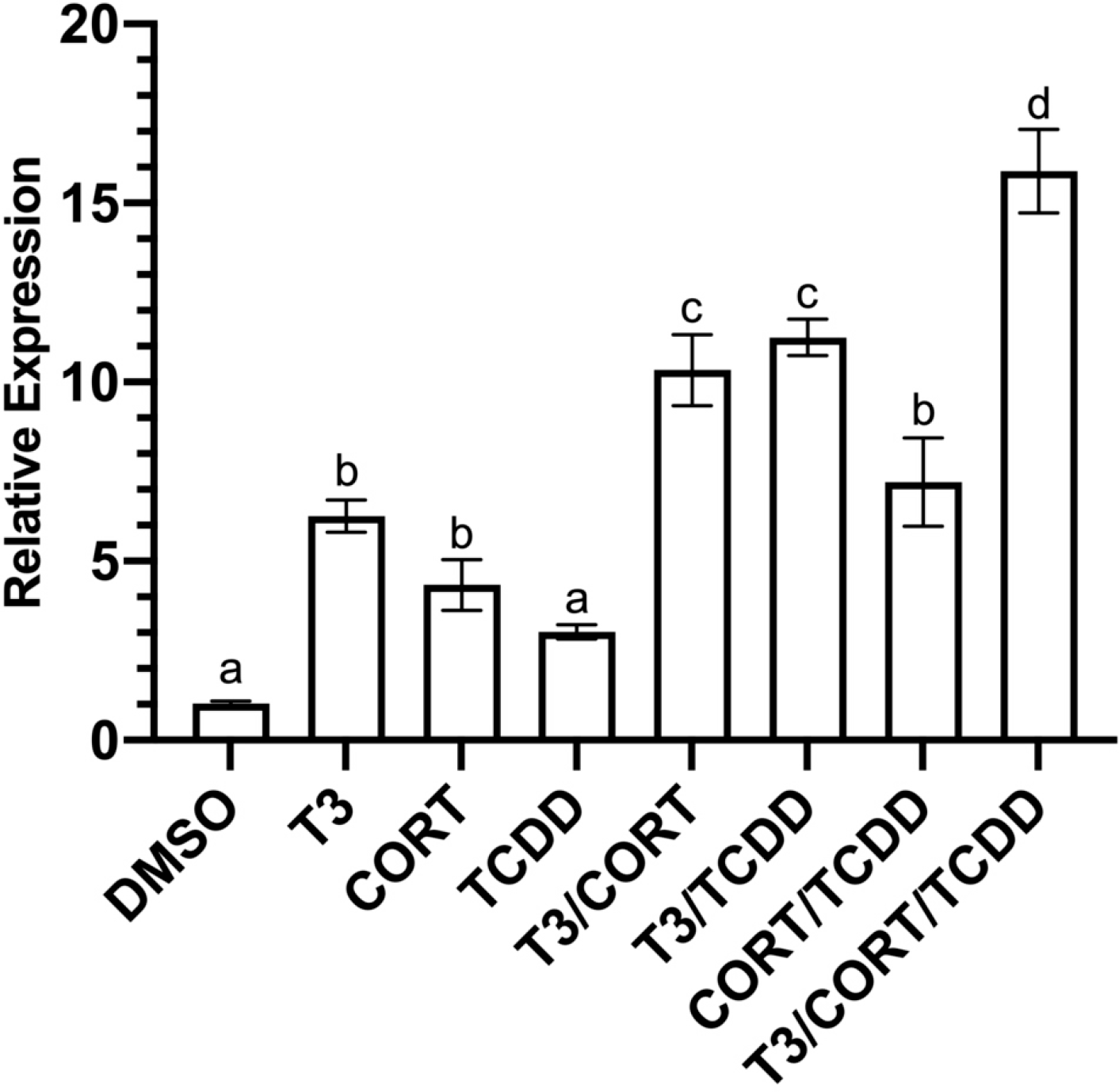
Induction of *klf9* mRNA in XLK-WG cells. Cells were incubated for 24 hours with 0.25% DMSO vehicle, 50 nM T3, 100 nM CORT, 175 nM TCDD, and all possible combinations in RPMI-1640 media supplemented with 20% charcoal-stripped FBS. Mean *klf9* mRNA induction relative to *actb* endogenous control was determined by RT- qPCR using the ΔΔCt method. *n* = 3 biological replicates per treatment group, 9 replicate measures in total. Error bars represent SEM. Statistical significance of differences between treatment groups were assessed by one-way ANOVA with Holm- Sidak’s multiple comparisons in GraphPad Prism 8.4.3. Values sharing a letter designation are not significantly different.

**Figure 2:**
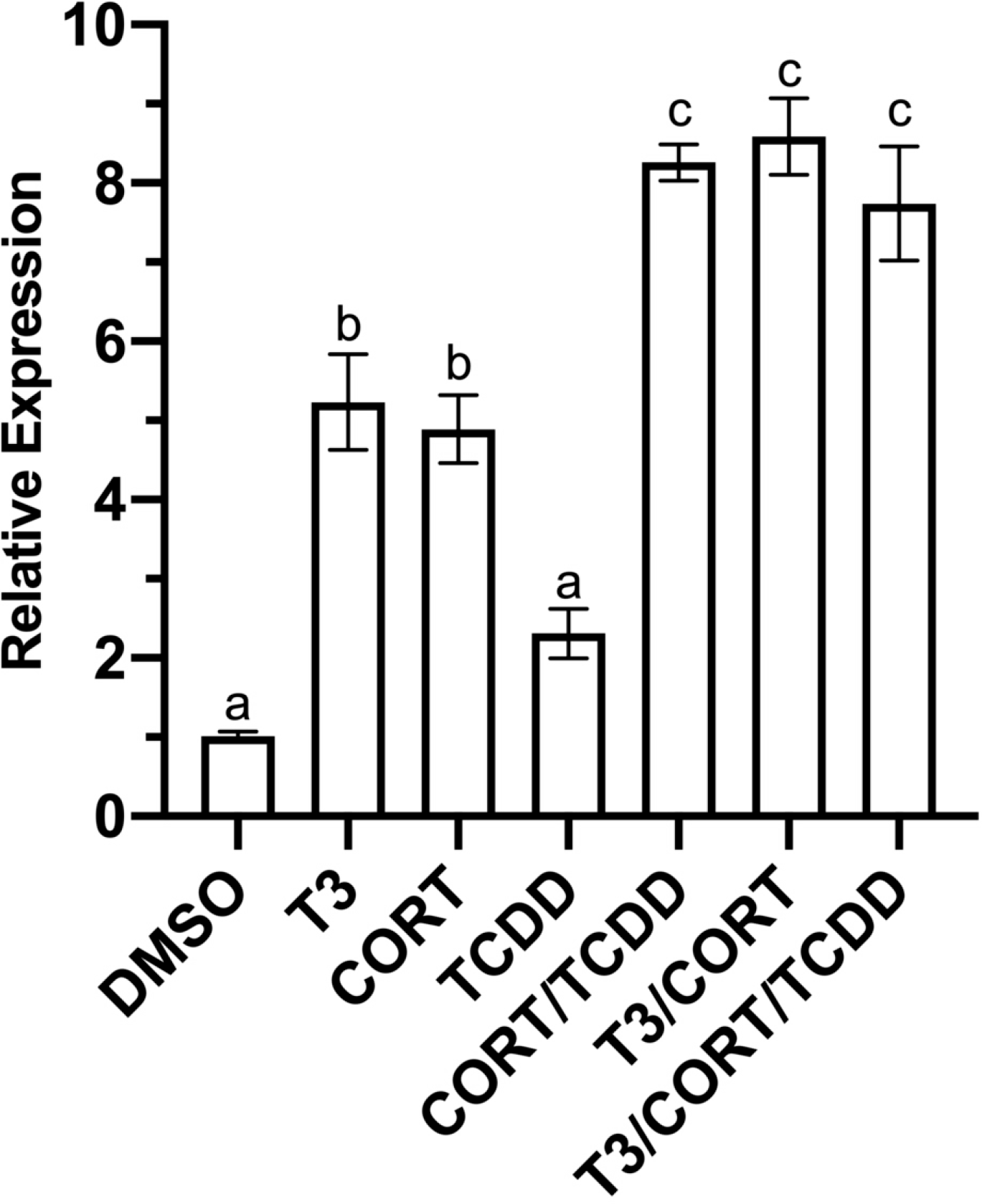
Induction of *klf9* mRNA in *X. laevis* tadpoles. Tadpoles (NF 52 to 54) were treated for 24 hours with 0.25% DMSO vehicle, 50 nM T3, 100 nM CORT, 175 nM TCDD, singly and in combination for 24 hours under a 12L:12D photoperiod. Mean *klf9* mRNA induction relative to *actb* endogenous control was determined by RT-qPCR using the ΔΔCt method. *n* = 3 tadpoles per treatment, 9 replicate measures in total. Error bars represent SEM. Statistical significance of differences between treatment groups and DMSO control were assessed by one-way ANOVA with Holm-Sidak’s multiple comparisons test in GraphPad Prism 8.4.3. Values sharing a letter designation are not significantly different.

### *klf9* Enhancer Analysis

To determine the molecular basis for the interactive effects of T3, CORT, and TCDD on *klf9* mRNA induction, we used segments of the upstream flanking region of the *Xenopus tropicalis klf9* promoter to conduct transactivation assays as described previously (Bagamasbad, et al., 2015). Locations of these segments relative to the TSS are denoted by their names (Fig. 3A). Each segment was cloned into pGL4.23 a firefly luciferase reporter vector. Reporter constructs bearing 1 kb segments up to -7 kb were a gift from Dr. Robert Denver (University of Michigan), while our group generated additional constructs to extend the range of analyzed sequences to -10 kb, deriving sequence information from the *X. tropicalis* genome browser (v9.2; Karimi, et al., 2018).

**Figure 3.**
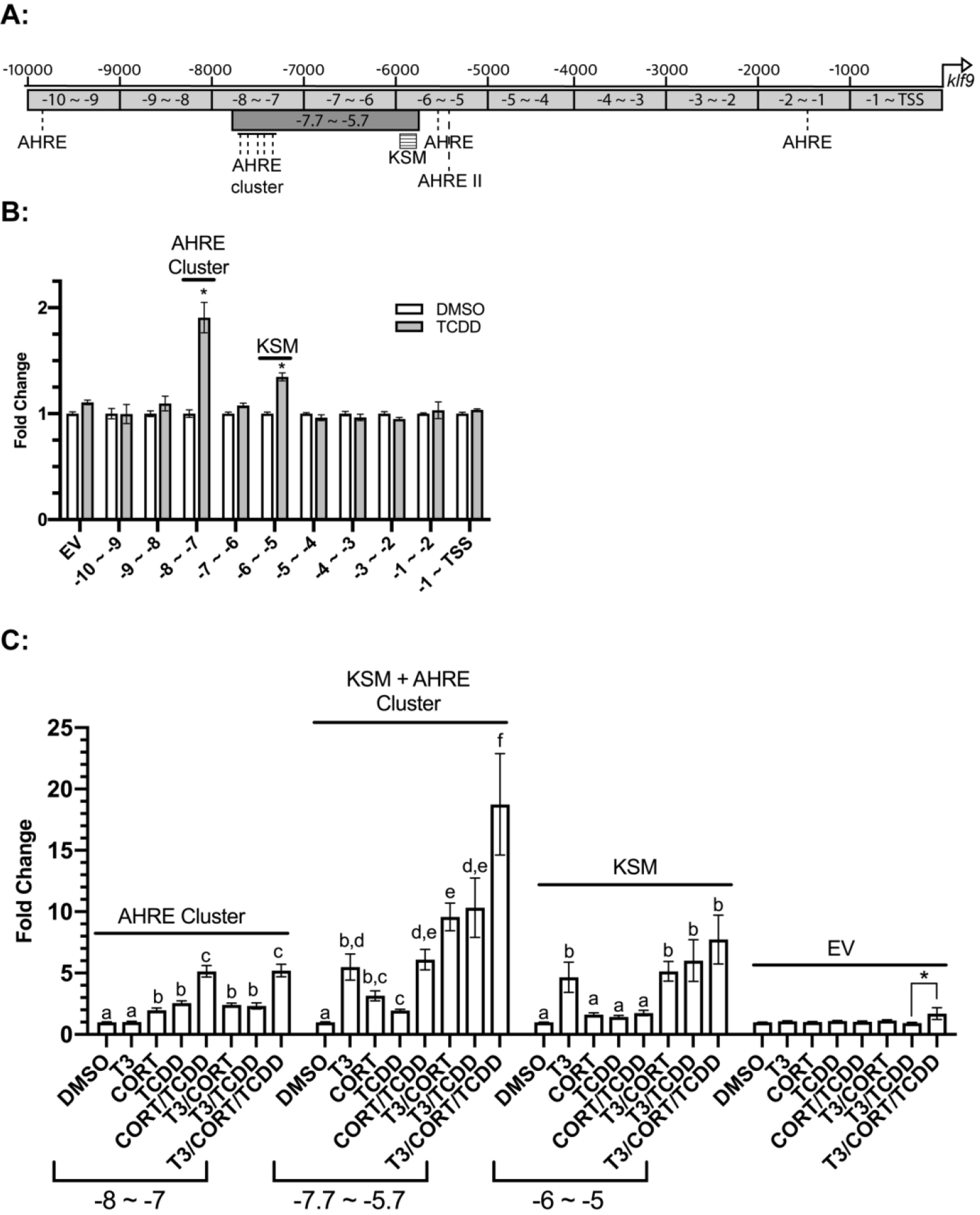
Reporter scan of three segments of the *X. tropicalis klf9* upstream region for CORT, T3, and TCDD responsiveness. **(A)** A map of relevant enhancers in the *klf9* upstream region. **(B)** Luciferase reporter gene expression induced by TCDD. **(C)** Luciferase reporter gene expression induced by T3, CORT, and/or TCDD. Transfected XLK-WG cells were incubated for 24 hours singly and in combination with DMSO vehicle, 50 nM T3, 100 nM CORT, 175 nM TCDD, in RPMI-1640 with 20% charcoal-stripped FBS. Reporter expression in lysates was measured by a dual luciferase assay. The EV construct lacks any *klf9* segment (“empty vector”). Values represent the mean fold change values of 3 independent biological replicates, each with three replicate measures. Error bars represent SEM. In Panel B, the statistical significance of response to TCDD exposure was assessed for each vector using student’s t-test. An asterisk (*) indicates significant difference from DMSO control. In Panel C, statistical significance of response to each treatment was assessed for each treatment by one-way ANOVA with Holm-Sidak’s multiple comparison test in GraphPad Prism 8.4.3, using log10-transformed values (Bagamasbad, et al., 2015). Values sharing a letter designation are not significantly different. An additional significant difference between treatments in the empty vector group is denoted with an asterisk (*).

We first sought to identify enhancers underlying elevated *klf9* mRNA expression by TCDD. We identified a cluster of five putative AHREs (Swanson, 2011) between -7 and -8 kb, as well as single AHREs in each of three segments located between -1 and - 2 kb, -5 and -6 kb, and -9 to -10 kb. A potential AHRE II element (Boutros *et al*., 2004; Sogawa *et al*., 2004) was also noted in the -5 to -6 kb region (Figure 3A). The newly discovered distal AHRE cluster (-7 ∼ -8) supported the strongest response to TCDD, a ∼2-fold increase, and appears to be the primary driver of AHR-mediated induction. In addition, the -6 to -5 kb construct drove reporter expression to a smaller degree (∼40% increase; Fig 3B). No other segments of the *klf9* upstream region supported TCDD-responsive luciferase induction, regardless of the presence of putative AHREs.

We next examined the interactive effects of T3, CORT, and TCDD on reporter gene expression in the most transcriptionally active regions. Consistent with studies published previously (Bagamasbad, et al., 2015), the response to T3 alone was driven exclusively by the *klf9* synergy module (KSM), located between -6 and -5 kb and downstream of the AHRE cluster (Fig. 3C and data not shown). The KSM also supported a small but statistically insignificant degree of responsiveness to CORT alone, while the -7 to -8 kb segment mediated a statistically significant response to CORT exposure (Fig. 3C).

To enable simultaneous assessment of both the KSM and the upstream AHRE cluster, an additional reporter plasmid was constructed to include both elements, encompassing the region between -5.7 and -7.7 kb (Fig. 3A). Simultaneous exposure to T3 and TCDD elicited a greater response than the CORT/TCDD co-exposure and rivaled the combination of T3 and CORT. The three agents together induced reporter expression over 18-fold, nearly double the value for the most efficacious combinations of two inducers (Fig. 3C).

### *klf9* upstream AHREs vary in strength

To investigate the functionality of each AHRE in the upstream cluster, we introduced a loss-of-function point mutation to each (5’CACGC3’ > 5’CATGC3’) in the context of the -5.7 to -7.7 kb construct. The AHREs are denoted numerically relative to the TSS, with the most distal as “AHRE 1” and the nearest “AHRE 5”. The plasmids carrying non-functional AHREs are named M1 through M5 to denote which AHRE is mutated. The M6 construct contains mutations in all 5 AHREs. Wholesale mutation of all five AHREs completely abrogated TCDD responsiveness (Fig. 4), indicating that the putative AHRE and AHREII located downstream of the KSM (see Fig. 3A) are not functional, at least in this heterologous assay. The M5 construct drove TCDD-induced reporter gene expression with comparably low efficacy to M6 (Fig. 4), suggesting that most transcriptional regulation of klf9 by AHR involves the AHRE on the downstream end of the cluster. Individual mutations of the remaining AHREs (constructs M1-M4) had insignificant effects on TCDD responsiveness (Fig. 4).

**Figure 4.**
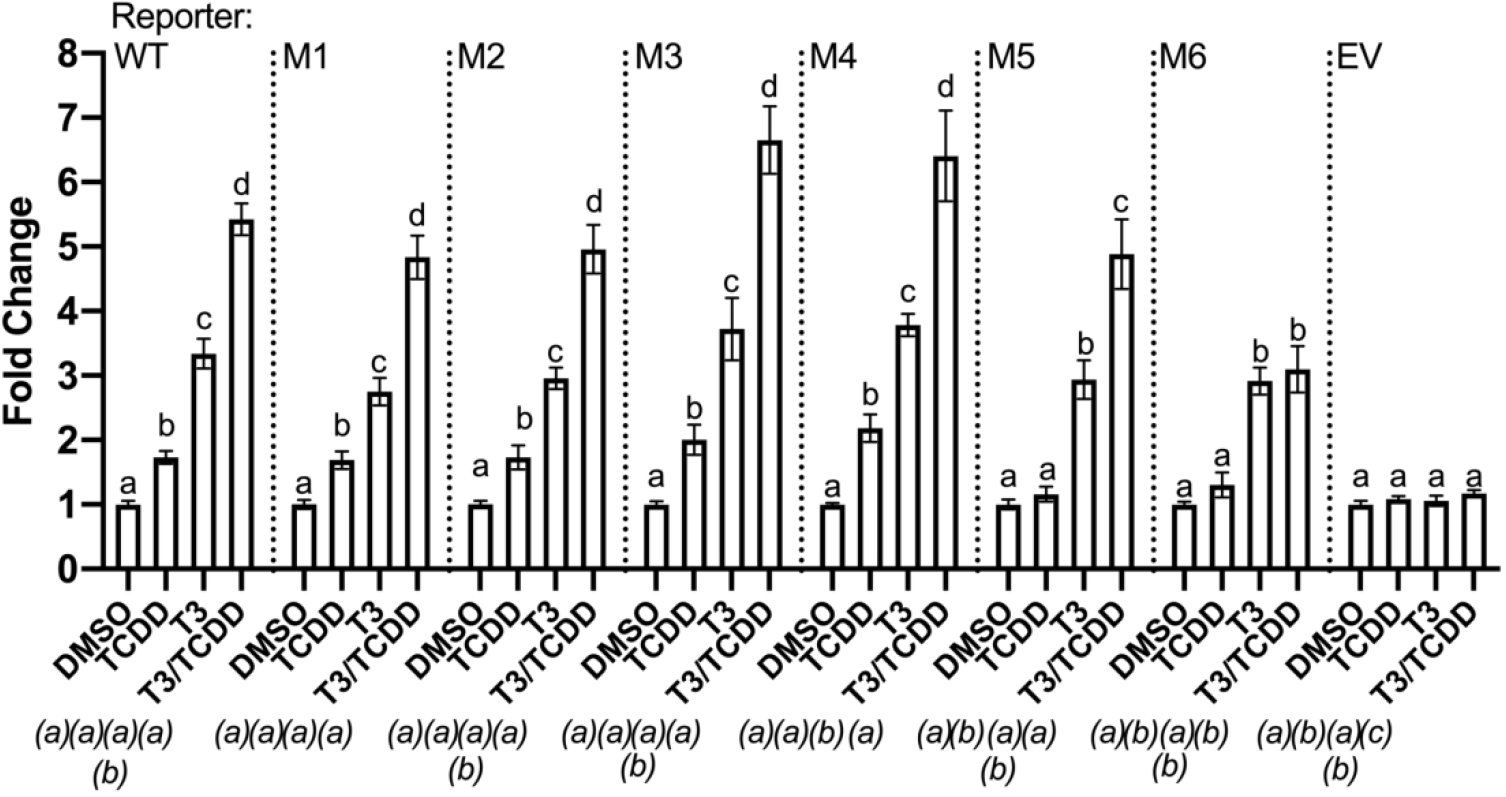
Functional assessment of individual AHREs within the predicted cluster upstream of *X. tropicalis klf9*. A transactivation assay compared the wild-type sequence and six AHRE mutant variants of the -5.7 to -7.7 kb segment of the *X. tropicalis klf9* upstream region cloned into pGL4.23 reporter vector. The five AHREs are numbered in order of their distance from the TSS, with the mutated AHRE farthest upstream denoted as “M1”. The M6 construct carries non-functional mutant versions of all 5 AHREs in the cluster. Transfected XLK-WG cells were treated for 24 hours with 0.25% DMSO vehicle, 50 nM T3, 175 nM TCDD, or a co-treatment of TCDD and T3 in RPMI-1640 + 20% FBS. Reporter expression in lysates was measured by a dual luciferase assay. The EV construct lacks any *klf9* segment (“empty vector”). Plotted values represent the mean fold change values of 3 independent biological replicates, each with three replicate measures. Error bars represent SEM. For each construct, statistical significance of response to each treatment was assessed for log10- transformed values (Bagamasbad, et al., 2015) by one-way ANOVA with Holm-Sidak’s multiple comparison test in GraphPad Prism 8.4.3. Values sharing a letter designation above each bar are not significantly different. Values for each treatment group were also statistically compared between vectors, with symbols to denote statistical significance appearing below each treatment group, italicized and in parentheses. Values sharing the same letter do not differ significantly from the same treatment in different reporter vectors.

### CORT responsiveness from a distal enhancer?

The -7 to -8 kb segment drove an unexpected response to CORT exposure in the absence of TCDD and a greater-than-additive response to treatment by both agents (Fig. 3C), suggesting the existence of a novel basis for *klf9* regulation by glucocorticoids outside the KSM. In a search of this region for conserved GRE sequences, we found no apparent local GR homodimeric binding sites on the sense strand in the 5’ to 3’ direction, nor any instance of the consensus GRE downstream half-site (TGTTCT), which may occur in combination with a second response element associated with a transcription factor able to complex with monomeric GR, such as XGRAF (Morin *et al*., 2000). However, we identified a previously-characterized GRE sequence, a GRE_CS8 (Okret *et al*., 1986), within the -7 to -8 kb region. To test the hypothesis that the sequence underlies CORT responsiveness, we generated a mutant -7 to -8 kb construct with the GRE_CS8 sequence, AGAACAGT, changed to TAGCATCT (Ren and Stiles, 1999). Compared with the wild-type construct, the GRE mutation conferred no observable decrease in the CORT-induced luciferase expression (Fig. 5), indicating that the consensus GRE-CS8 is not directly involved with CORT signaling.

**Figure 5.**
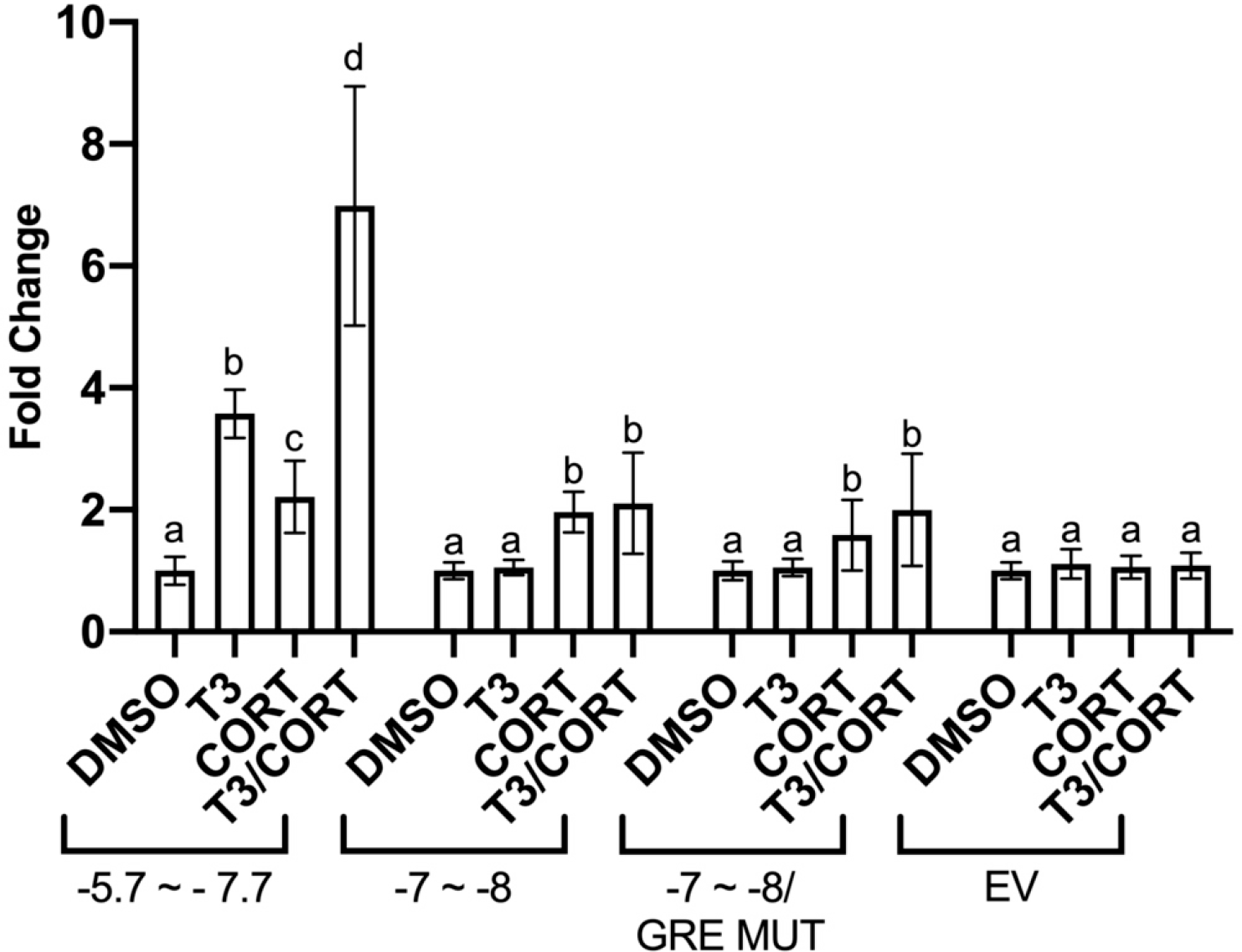
Functional assessment of a candidate GRE. Transactivation assay examined CORT responsiveness of indicated *X. tropicalis klf9* upstream regions. The GRE MUT construct contains the -7 to -8 kb region in which the putative GRE contains a mutation to block GR binding. The -5.7 to -7.7 kb segment (a positive control for CORT responsiveness) includes the KSM but lacks the upstream putative GRE site. Transfected XLK-WG cells were incubated for 24 hours with 0.25% DMSO vehicle, 50 nM T_3_, 100 nM CORT, or a co-treatment in RPMI-1640 with 20% charcoal-stripped FBS. Reporter expression in lysates was measured by a dual luciferase assay. Bars represent the mean fold change values of 3 independent biological replicates, each with three replicate measures. Error bars represent SEM. For each construct, statistical significance of response to each treatment was assessed for log10-transformed values (Bagamasbad, et al., 2015) by one-way ANOVA with Holm-Sidak’s multiple comparison test in GraphPad Prism 8.4.3. Values sharing a letter designation are not significantly different.

### Conservation of frog *klf9* enhancers in humans

Frog metamorphosis represents a widely used model for the study of the endocrinology and toxicology of human development, especially relating to thyroid hormone (Buchholz, 2015). To establish the human health relevance of TCDD’s disruption of endocrine regulation of *klf9* expression in tadpoles, we sought to investigate the degree of conservation of the sequence and function of the *klf9* upstream region shared between these species. We first tested the hypothesis that TCDD alters T3 induction of *KLF9*, as it does in the frog system (Figs 1-3 and Taft, et al., 2018). mRNA abundance was measured in HepG2 cells (human liver hepatocellular carcinoma), a widely used toxicological model in the study of xenobiotic metabolism, including the role of AHR. HepG2 cells exhibited clear *KLF9* mRNA induction by T3 (100 nM) and TCDD (10 nM), and co-treatment with TCDD boosted the response nearly 35% (Fig. 6). Notably, TCDD alone induced mean *CYP1A1* expression by ∼700-fold, demonstrating the efficacy of this concentration in these assays (Fig. 6B).

**Figure 6.**
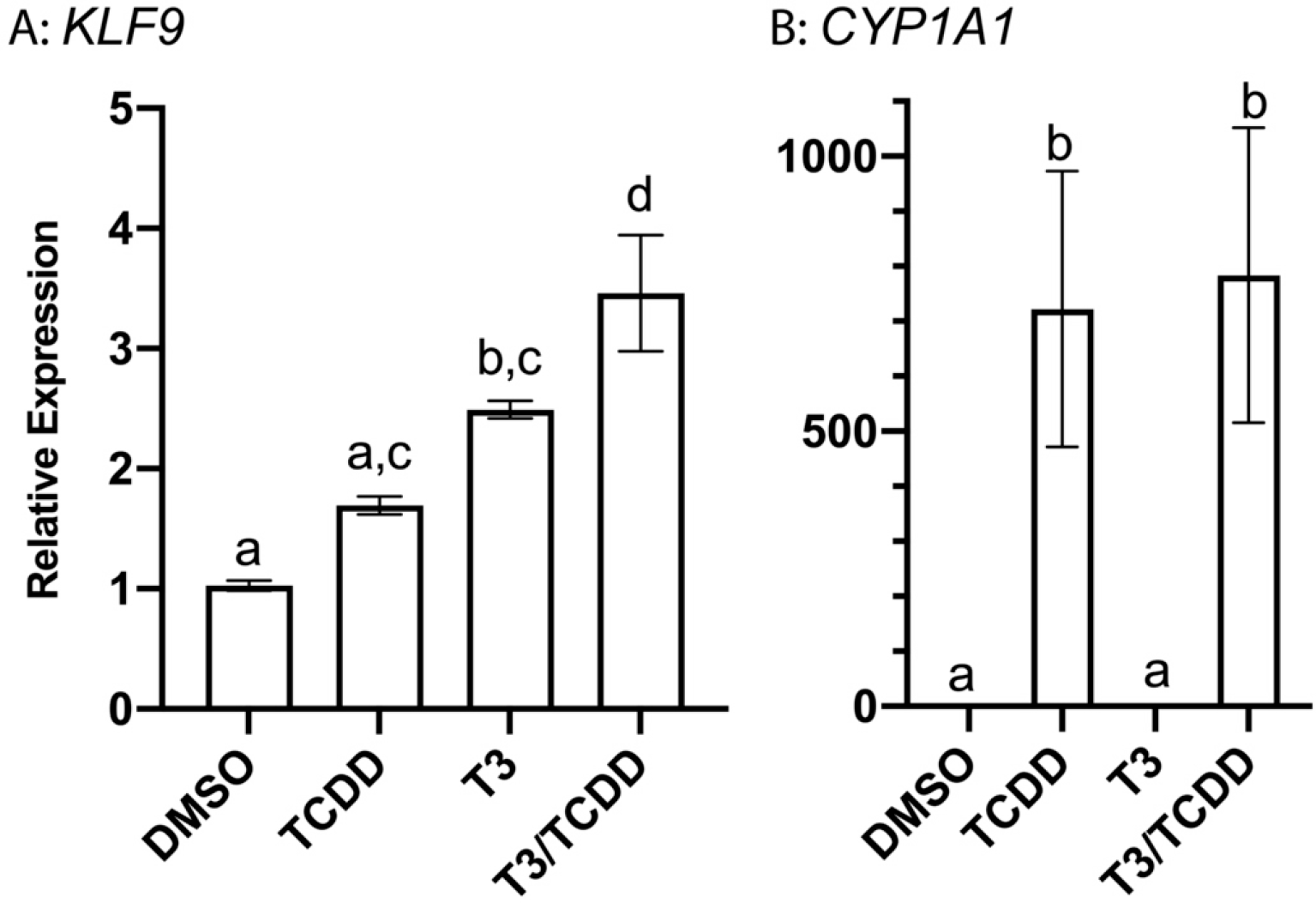
Induction of *KLF9* mRNA in HepG2 cells. Cells were incubated for 24 hours with 0.25% DMSO vehicle, 10 nM TCDD, 100 nM T_3_, or a co-treatment in EMEM supplemented with 10% charcoal-stripped FBS. Mean *KLF9* induction (panel A) was calculated relative to *GAPDH* endogenous control by RT-qPCR using the ΔΔCt method. *CYP1A1* induction (panel B) was also included as a positive control of TCDD responsiveness. *n* = 3 biological replicates per treatment group, each with three replicate measures. Error bars represent SEM. Statistical significance of differences between treatment groups were assessed by one-way ANOVA with a Holm-Sidak’s multiple comparisons test in GraphPad Prism 8.4.3. Values sharing a letter designation are not significantly different.

We next explored the organization of enhancers driving human *KLF9* expression, revealing a pattern associated with the human gene that is strongly analogous to its counterpart in the frog genome. TRE and GRE elements are situated within the highly conserved KSM (Bagamasbad, et al., 2015), ∼4.5 kb upstream to the promoter. In addition, a cluster of 6 core AHREs lies upstream of the KSM, although much closer (less than 1 kb) than in the frog genome. Additional solitary AHREs flank the KSM in both nearby and distant positions (Fig. 7A). To assess response element functionality, we next performed luciferase-reporter assays in HepG2 cells, examining the response to T3 and TCDD. Reporter constructs contained either the 6-AHRE cluster (-4.7 to -5.4 kb), the KSM (-4.0 to -4.7 kb), or both groups of enhancers (-4.0 to -5.4 kb). Again, the induction pattern bore striking resemblance to that seen in the frog cells (Fig. 7B). The AHRE cluster (-5.4 to -4.7 kb) supported only TCDD responsiveness. The KSM reporter (-4.7 to -4 kb) was induced 2-fold by T3, but TCDD induced no detectable response and did not potentiate T3-driven transactivation. The reporter containing both the KSM and the AHRE cluster was responsive to both agents and exhibited a greater T3 response than did the shorter KSM construct. We conclude that in at least in HepG2 cells, TCDD can disrupt T3-induced *KLF9* through a mechanism parallel to that of the frog system.

**Figure 7.**
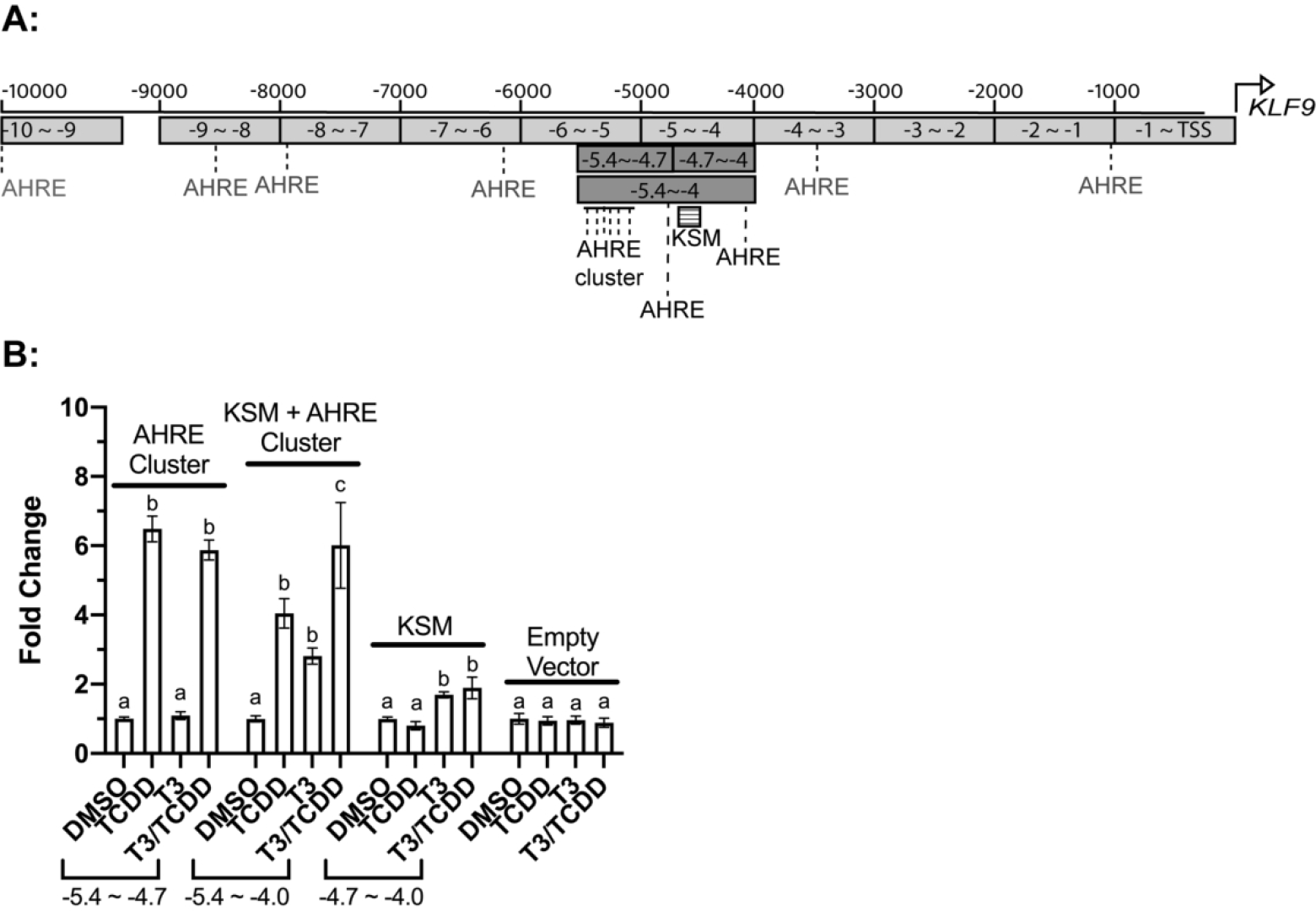
Enhancer analysis of three segments of the human *KLF9* upstream region. Transfected HepG2 cells were incubated for 24 hours with 0.25% DMSO vehicle, 10 nM TCDD, 100 nM T_3_, or a co-treatment in EMEM supplemented with 10% charcoal-stripped FBS. Reporter expression in lysates was measured by a dual luciferase assay. Bars represent the mean fold change values of 3 independent biological replicates, each with three repeated measures. Error bars represent SEM. For each construct, statistical significance of response to each treatment was assessed for log10-transformed values (Bagamasbad, et al., 2015) by one-way ANOVA with Holm- Sidak’s multiple comparison test in GraphPad Prism 8.4.3. Values sharing a letter designation are not significantly different.

## Discussion

*Xenopus* metamorphosis represents a widely used model system for studying post-embryonic development, including the disruption of endocrine regulation by xenobiotics (EPA, 2011). Thyroid hormone plays a central role in the activation and repression of gene expression that directs the metamorphosis of tadpoles into frogs (Buchholz, et al., 2006; Sachs *et al*., 2000; Shi, 2009), and expression of either TRα or TRβ is required for survival beyond the metamorphic climax (NF 61; Shibata *et al*., 2020). Glucocorticoids represent an ancillary endocrine influence on metamorphosis, regulating the rate at which it proceeds by altering sensitivity of target tissues to TH and modulating the expression of TH-regulated genes (Sachs and Buchholz, 2019). The significance of the role of GCs is underscored by the requirement of both CORT and GR for survival and completion of metamorphosis in *Xenopus* (Shewade *et al*., 2020; Sterner *et al*., 2020). The endocrine balance of this system is subject to toxicological disruption by dioxin-like chemicals. Expression of several TR target genes can be induced by TCDD and other AHR agonists, and TCDD exposure can alter tail resorption and limb growth during metamorphosis (Taft, et al., 2018). In this study we sought to characterize upstream enhancers that underlie the AHR contribution to the regulation of *klf9*, an immediate early gene in the cascade of expression changes in the TH- and GC-mediated control of metamorphosis (Bonett *et al*., 2010; Bonett *et al*., 2009; Furlow and Kanamori, 2002). We demonstrate the importance of a novel upstream cluster of AHREs in the regulation of *klf9* mRNA expression. Conservation of this AHRE cluster in the human genome suggests its importance in *klf9*-mediated developmental and physiological processes common to people and frogs as well as the potential for cross- talk with and disruption of TR and GR activity.

### Glucocorticoid regulation of *Xenopus klf9*

This toxicological study extends the fundamental understanding of endocrine control of *klf9* induction by glucocorticoid receptors. While the role of TR and GR binding within the *klf9* synergy module is a well-documented requirement for *klf9* induction (Bagamasbad, et al., 2015), our results point to a novel role for GR outside the KSM. Combined CORT/TCDD exposure drove transactivation associated with two *klf9* reporter constructs—the -7 to -8 kb region (containing the AHRE cluster) and the -5.7 to -7.7 kb region (AHRE cluster + KSM)—but not -5 to -6 kb region (KSM alone; Fig. 3). Thus, the functional interaction of AHR and GR does not require the well-characterized GRE in the KSM that drives synergistic *klf9* induction in conjunction with TR (Bagamasbad, et al., 2015). Examining a possible alternative, we tested the functionality of the only readily predicted GRE motif in the -7 to -8 kb region (GRE_CS8, AGAACAGT; (Okret, et al., 1986), showing that CORT-driven transactivation was unchanged following mutation of this site. This result is not unreasonable. While rat GR can bind the GRE_CS8 sequence (Okret, et al., 1986), mutation of the motif in the upstream region of the human *A_1_AR* gene failed to reduce dexamethasone-stimulated transactivation (Ren and Stiles, 1999). *In silico* identification of functional GREs from nucleotide sequence alone presents a complex challenge. GR binds only a fraction of predicted response elements, and those vary with cell type and physiological context. Functional GREs can be located far from the core promoter, and GR can interact with poorly conserved, non-classical response elements (Burd and Archer, 2013). The most predictively useful characteristics of functional GREs include chromatin accessibility (Burd and Archer, 2013), evolutionary conservation (So *et al*., 2008), and the presence of proximal binding sites for additional transcription factors (Datson *et al*., 2011).

GR is also known to regulate transcription of certain targets via protein:protein interactions with DNA-binding partners like AP-1 and NF-1 (Scheschowitsch, et al., 2017). *In vivo*, many key developmental events do not appear to rely on direct GR:DNA binding. Although GR expression is necessary for survival in mice, mouse strains expressing mutant GR lacking the ability to form the canonical DNA-binding homodimer nonetheless survived birth (Reichardt and Schutz, 1998). The ability of activated AHR and GR to interact directly has been demonstrated by immunoprecipitation, and AHR activation by 3-methylcholanthrene coupled with GR activation by dexamethasone led to recruitment of AHR to a GRE and a synergistic increase in metallothionein 2A (*MT2A*) mRNA in HeLa cells (Sato *et al*., 2013). Furthermore, increased transactivation driven by a GR:AHR complex has been observed in GRE-driven reporter gene assays in human cell lines stimulated with dexamethasone and the AHR agonist benzo[*a*]pyrene (Wang *et al*., 2009). These effects are gene-specific, as cotreatment with dexamethasone and TCDD has been found to increase the recruitment of GR and AHR to select genes in ARPE-19 cells (*e.g., ANGPLT4* and *FKBP5*) but to limit AHR association with response elements associated with other genes, including the canonical AHR targets *CYP1A1*, *CYP1B1*, and *AHRR* (Jin *et al*., 2017). Similar protein:protein interactions may underlie the CORT-induced transactivation driven by the -7 to -8 kb region that lacks an apparent functional GRE motif.

Our examination of the -5 to -6 kb region reinforces previously published evidence that activated GR directly binds the conserved classical GRE in the KSM (Bagamasbad, et al., 2015; Mostafa *et al*., 2021), though the magnitude of our observed CORT response is lower than values reported previously (Bagamasbad, et al., 2015). We inconsistently observed transactivation induced solely by CORT in some individual experiments (data not shown), but the largest and most consistent CORT response occurred with the -5.7 to -7.7 kb reporter construct, which includes the KSM but may also overlap with the upstream CORT-responsive segment (-7 to -8 kb). Thus, the KSM GRE alone may not be sufficient for maximal CORT induction of *klf9*. The conservation of this additional functional GRE locus in humans is not yet clear.

### Enhancers driving disruption of *klf9* expression by TCDD

These experiments identify a cluster of functional AHREs upstream of the *X. tropicalis klf9* synergy module. This arrangement of enhancers is mirrored by a strong TCDD response proximal to the human KSM in a region containing several predicted AHREs. Mutation of individual AHREs within the *X. tropicalis* cluster was associated with varying degrees of response to TCDD, while the simultaneous mutation of all five AHREs abrogated the TCDD response and the additive CORT/TCDD response (Fig. 4). These transactivation results are consistent with reports of AHR and AHR Repressor ChIPseq peaks upstream of human *KLF9* in MCF-7 cells (Yang *et al*., 2018 and corresponding data in the GEO, GSE 90550). The position of these peaks (-5371) directly corresponds to the cluster of functional AHREs we characterized in HepG2 cells. AHRE clusters have also been identified in the regulatory region of other dioxin-induced genes, most notably mouse *Cyp1a1* and *Cyp1a2* (Nukaya *et al*., 2009). The variable degree of functionality of individual AHREs in the *X. tropicalis* cluster (Fig. 4) appears to be common to AHRE clusters in other genes, including mouse *Cyp1a2* (Kawasaki *et al*., 2010) and zebrafish *cyp1a* (Zeruth and Pollenz, 2007).

In experiments comparing transactivation driven by combined T3/CORT/TCDD exposure associated with plasmids containing the -5.7 to -7.7 kb region and those containing the -5 to -6 kb region, the longer segment supported a greater response (Fig. 3C). This effect is likely driven by increased TR and GR binding at the KSM occurring in concert with AHR-driven transcriptional upregulation linked to the upstream AHRE cluster. This discrepancy may also result from differences in plasmid structure. The plasmids used in transactivation assays are necessarily imperfect models of gene regulation, as they lack the capacity for long-range transcription factor interactions and undergo dynamic conformational changes in ways distinct from genomic DNA (Higgins and Vologodskii, 2015). Likewise, altered chromatin accessibility driven by nucleosome alterations, a key aspect of transcription factor function *in vivo*, may not occur uniformly in all plasmid models. While nucleosome formation has been reported from supercoiled plasmid DNA (Nakagawa *et al*., 2001), total plasmid size is a factor in chromatin formation (Jeong *et al*., 1991). Regardless, previous observations that ligand-bound TR can drive transcription irrespective of promoter chromatinization (Shi, 2000) provide a basis for our observation the that T_3_-driven transactivation response associated with the -5 to -6 kb region plasmid was of a similar magnitude to that of the -5.7 to -7.7 kb region plasmid while the GR and AHR responses varied more substantially. A full accounting of the mechanisms by which these nuclear receptors interact to alter transcription will require chromatin immunoprecipitation of factors bound within the actual genome, not merely a heterologous system. These experiments, including the development of antibodies amenable to use with the frog orthologs, will be the subject of future studies.

## Conclusions and Toxicological Implications

Building on previous experiments that identified the *klf9* synergy module (Bagamasbad, et al., 2015), this work provides a molecular basis for dioxin-driven disruption of endocrine-mediated *klf9* induction in frogs. It also identifies a second locus of CORT response. We propose an updated model of *klf9* regulation by TRs, GR, and AHR (Fig. 8). This model includes a CORT-responsive site and an AHRE cluster between -7 and -8 kb that act in concert with the established roles of the evolutionarily conserved GRE and TRE in the KSM. Additionally, two potential sources of the modest AHR transactivation mediated by elements proximal to the KSM itself are indicated, which include tethering of the AHR-ARNT complex with DNA-bound transcription factors within the KSM and direct DNA binding at putative AHREs. We propose that the activated TR and GR pathways interact to regulate *klf9* induction, and that TCDD can disrupt this process by further increasing *klf9* expression through “cross-talk” between AHR and the two hormone receptors, disrupting baseline endocrine control. These findings provide new and promising avenues for the study of the TR/GR/AHR regulatory axis not only in *Xenopus* but also in animal models for developmental toxicology across the tetrapod lineage. They expand the focus on enhancers regulating *klf9* transcriptional regulation beyond the well-studied *klf9* synergy module (Bagamasbad et al. 2015). They also define an important early event—binding of AHR to the distal AHRE cluster—in the emerging adverse outcomes pathway underlying the effects of TCDD on limb growth and tail resorption during *Xenopus* metamorphosis as well as events in human development and physiology that are under control of TR, GR, and *klf9*.

**Figure 8.**
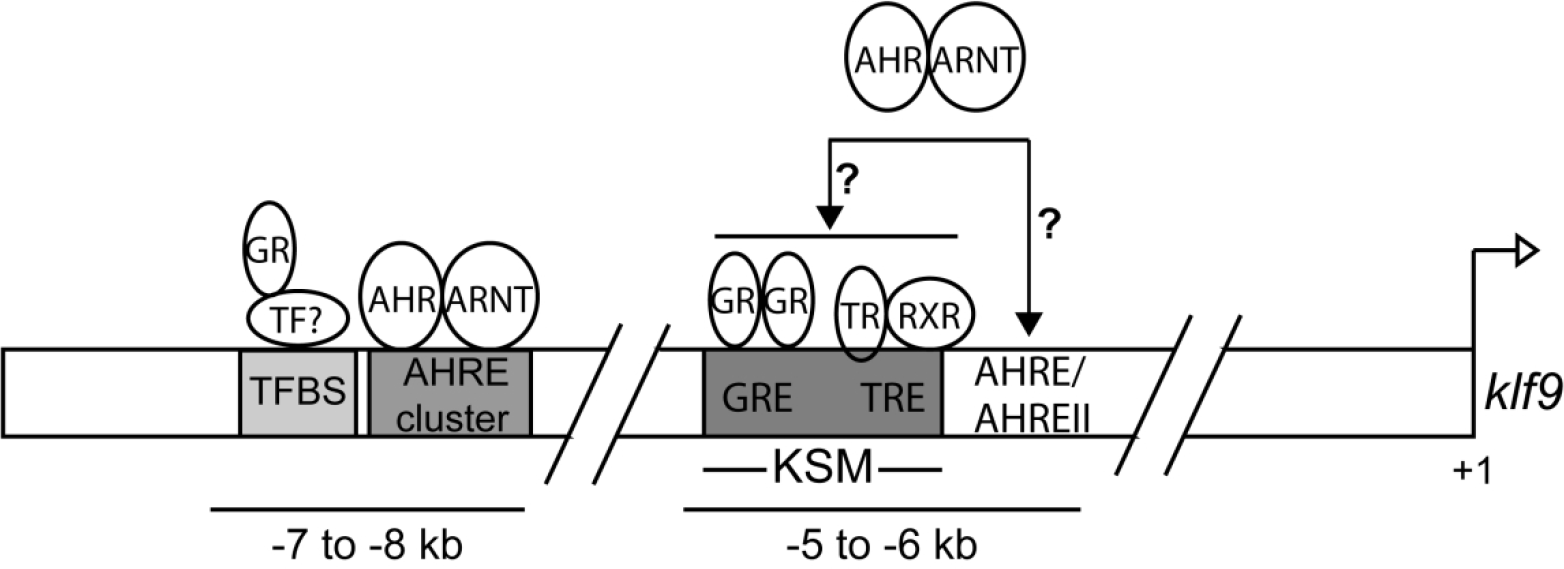
A model for frog *klf9* regulation by T3, CORT, and TCDD. Enhancer elements demonstrated to be engaged by specific receptors are indicated. Hypothetical interactions are denoted by a question mark. TF, Transcription factor; TFBS, Transcription Factor Binding Site.

## Acknowledgements

We gratefully acknowledge Dr. Pia Bagamasbad (University of Michigan) for generating multiple reporter gene constructs for transactivation assays and Prof. Robert J. Denver (University of Michigan) for making them available for our studies.

